# Neurons in the dorsomedial hypothalamus promote, prolong, and deepen torpor in the mouse

**DOI:** 10.1101/2021.09.05.458994

**Authors:** Michael Ambler, Timna Hitrec, Andrew Wilson, Matteo Cerri, Anthony Pickering

## Abstract

Torpor is a naturally occurring, hypometabolic, hypothermic state engaged by a wide range of animals in response to imbalance between the supply and demand for nutrients. Recent work has identified some of the key neuronal populations involved in daily torpor induction in mice, in particular projections from the preoptic area of the hypothalamus (POA) to the dorsomedial hypothalamus (DMH). The DMH plays a role in thermoregulation, control of energy expenditure, and circadian rhythms, making it well positioned to contribute to the expression of torpor. We used activity dependent genetic TRAPing techniques to target DMH neurons that were active during natural torpor bouts in female mice. Chemogenetic reactivation of torpor-TRAPed DMH neurons in calorie-restricted mice promoted torpor, resulting in longer and deeper torpor bouts. Chemogenetic inhibition of torpor-TRAPed DMH neurons did not block torpor entry, suggesting a modulatory role for the DMH in the control of torpor. This work adds to the evidence that the POA and the DMH form part of a circuit within the mouse hypothalamus that controls entry into daily torpor.

**Significance:** Daily heterotherms such as mice employ torpor to cope with environments in which the supply of metabolic fuel is not sufficient for the maintenance of normothermia. Daily torpor involves reductions in body temperature, as well as active suppression of heart rate and metabolism. How the central nervous system controls this profound deviation from normal homeostasis is not known, but a projection from the preoptic area to the dorsomedial hypothalamus has recently been implicated. We demonstrate that the dorsomedial hypothalamus contains neurons that are active during torpor. Activity in these neurons promotes torpor entry and maintenance, but their activation alone does not appear to be sufficient for torpor entry.

## Introduction

Torpor is the naturally occurring hypothermic, hypometabolic, and hypoactive component of hibernation, which can be prolonged (in seasonal hibernators), or brief (in daily heterotherms such as the mouse). It serves as an adaptive response to relative energy deficit: a controlled reduction in metabolic demand in response to reduced availability of substrate. During torpor, body temperature typically runs a few degrees above ambient temperature, which in hibernating arctic ground squirrels results in core temperatures as low as −2.9°C (Barnes, 1989). Metabolic rate falls to between 1 and 5% of euthermic rates with similar reductions in heart and respiratory rates (Heldmaier et al., 2004).

The mechanisms controlling this profound deviation from normal physiology are not known, but recent work has indicated a role for the preoptic area of the hypothalamus (POA) in generating a hypothermic, hypometabolic, bradycardic state similar to daily torpor in the mouse (Hrvatin et al., 2020; Takahashi et al., 2020; Zhang et al., 2020). These neurons are active during natural fasting-induced daily torpor (Hrvatin et al., 2020; Zhang et al., 2020); are necessary for the full expression of daily torpor (Hrvatin et al., 2020; Takahashi et al., 2020; Zhang et al., 2020); express *Adcyap1* (Hrvatin et al., 2020), and/or pyroglutamylated RFamide peptide (QRFP) (Takahashi et al., 2020), and/or oestrogen receptors (Zhang et al., 2020), with evidence of overlap in these markers. These studies suggest that activation of a projection from the POA to the dorsomedial hypothalamus (DMH) contributes to the generation of this torpor-like state, although projections from the POA elsewhere, for example to arcuate, may also contribute (Hrvatin et al., 2020; Zhang et al., 2020).

Projections from the POA to DMH are also involved in thermoregulation outside of the context of torpor. Warm-activated neurons in the ventral part of the lateral POA (vLPO) project to the DMH and inhibit thermogenesis and locomotion (Zhao et al., 2017). Similarly, warm-activated neurons in the ventromedial POA (VMPO) that express *Adcyap1* and brain-derived neurotrophic factor (BDNF) project to the DMH and suppress thermogenesis (Tan et al., 2016). These thermogenesis inhibiting warm-activated POA to DMH projections that play a role in thermoregulation are GABAergic (Tan et al., 2016; Zhao et al., 2017), in contrast to the predominantly glutamatergic projections that are implicated in torpor (Hrvatin et al., 2020; Takahashi et al., 2020). In addition to an established role in thermoregulation, the DMH also regulates energy balance and food intake (Bi et al., 2012; Jeong et al., 2015; Yang et al., 2009), heart rate (Piñol et al., 2018), and plays a role in adjusting circadian rhythms in response to food availability (Chou et al., 2003; Gooley et al., 2006; Mieda et al., 2006). Hence, the DMH is well positioned to play a role in the regulation of torpor, and there is evidence to support its activation during daily torpor in the mouse (Hitrec et al., 2019).

We hypothesised that the DMH plays a role in the control of torpor. To test this hypothesis we used activity-dependent targeting of neurons (DeNardo et al., 2019) within the DMH that were active during natural torpor bouts in female mice. Using this approach, we first labelled neurons within the hypothalamus that were active during torpor. Next, we expressed excitatory or inhibitory DREADDs (Roth, 2016) in DMH neurons that were active during torpor, and evaluated the effects of activating these receptors on the propensity to enter torpor during calorie restriction.

## Materials & Methods

### Mice

All studies had the approval of the local University of Bristol Animal Welfare and Ethical Review Board and were carried out in accordance with the UK Animals (Scientific Procedures) Act. Female TRAP2 (Allen et al., 2017; DeNardo et al., 2019), Ai14 (Madisen et al., 2010), and C57BL/6J mice, at least 8 weeks of age with mean body weight 22.9 ± 2.8 grams on experiment entry were used. Female mice were chosen because of their increased propensity to undergo torpor (Swoap and Gutilla, 2009). Mice were maintained on a reversed 12-hour light/dark cycle with lights off at 08:30, hence lights off was assigned as zeitgeber time (ZT) 0. Ambient temperature was 20.9 ± 0.4°C. Mice had free access to water and standard mouse chow (LabDiet, St. Louis, MO 63144, USA) except during periods of calorie restriction when food was limited as detailed below. They were housed in groups of up to four.

A homozygous breeding colony was established from heterozygous TRAP2 mice kindly donated by the Liqun Luo laboratory in Stanford University, California. The strain is now available via Jackson laboratory (www.jax.org/strain/030323). The TRAP2 mouse line carries the gene for Cre-ERT2 under the fos promoter leading to Cre-ERT2 expression in active neurons. Homozygous Ai14 mice were obtained from the University of Bristol in-house colony, having been originally purchased from Jackson laboratories (www.jax.org/strain/007908). The Ai14 mouse carries a floxed gene encoding a red fluorophore (tdTomato) knocked into the *Gt(ROSA)26Sor* locus. This requires the action of Cre to remove a stop codon to allow tdTomato expression. Breeding pairs were established with homozygous TRAP2 and homozygous Ai14 mice to generate TRAP:Ai14 double-heterozygous offspring. The resultant double-transgenic TRAP:Ai14 mice produce red fluorescent tdTomato protein in neurons that were active during the time period defined by the injection of 4-hydroxytamoxifen (4-OHT).

### Viral vectors

Three viral vectors were used:

AAV2-hSyn-DIO-hM3Dq-mCherry was a gift from Bryan Roth (Addgene viral prep #44361-AAV2; www.addgene.org/44361) (Krashes et al., 2011); 4.6×10^12^ viral genome copies per ml. This vector delivered a Cre-dependent mCherry-tagged excitatory DREADD gene under the human synapsin promoter. It was mixed in a 4:1 ratio with the EGFP-expressing vector (used to identify the injection site), giving a final titre for this vector of 3.7×10^12^ viral genomes per ml.

AAV2-hSyn-DIO-hM4Di-mCherry was a gift from Bryan Roth (Addgene viral prep #44362-AAV2; www.addgene.org/44362) (Krashes et al., 2011); 2×10^13^ viral genome copies per ml. This vector delivered a Cre-dependent mCherry-tagged inhibitory DREADD gene under the human synapsin promoter. It was mixed in a 1:2 ratio with the EGFP-expressing vector, and the resulting vector mixture further diluted in a 10-fold with sterile phosphate buffered saline, giving a final titre of 6.7×10^11^ viral genomes per ml.

pAAV2-CMV-PI-EGFP-WPRE-bGH was a gift from James M. Wilson (Addgene viral prep #105530-AAV2; www.addgene.org/105530); 7×10^12^ viral genome copies per ml. This vector delivered the gene coding for enhanced green fluorescent protein (EGFP) under the ubiquitous CMV promoter. This vector was used to confirm the localisation of injection, because the expression of mCherry fluorescence in the two vectors above is contingent on successful TRAPing (and so would not be visible if the injected area is not TRAPed).

### Stereotaxic injection of viral vectors

Mice were anaesthetised with ketamine (70mg/kg i.p.) and medetomidine (0.5mg/kg i.p.). Depth of anaesthesia was assessed and monitored by loss of hind paw withdrawal reflex. Additional i.p injections of anaesthetic were administered as needed to maintain surgical depth of anaesthesia. Core temperature was maintained using a servo-controlled heat pad and a rectal temperature probe (Harvard Apparatus). The planned incision site was shaved, and skin cleaned with iodine solution and sterile surgical technique was used throughout. Anaesthetised mice were placed in a stereotaxic frame, the head was fixed in atraumatic ear bars and skull position maintained horizontal by an incisor bar (David Kopf Instruments, USA).

Microcapillary pipettes were made from microcapillary glass (Sigma, USA) on a vertical pipette puller (Harvard Apparatus, UK). Pipettes were filled with mineral oil then vector was back-filled using a robotic microinjector (Nano-W wireless capillary microinjector, Neurostar, Germany), producing a visible vector-mineral oil interface. The scalp was incised in the midline and burr holes made bilaterally at bregma −1.8mm, lateral ±1mm with a drill attachment (Neurostar, Germany). The microcapillary pipette was inserted at an angle of 8° towards the midline. Bilateral injections were made at depths of 5 and 4.75mm relative to the surface of the brain. Each injection was 180nl and was delivered at a rate of 100nl/minute. The injection pipette remained in place for one minute after the first injection and for five minutes after the second before removing.

Following vector injections, the wound was closed with non-absorbable suture and dressed with antibacterial wound powder. Anaesthesia was reversed with atipamezole (1mg/kg i.p, Antisedan, Zoetis). Buprenorphine was administered for analgesia (0.1mg/kg s.c., Vetergesic, Ceva Animal Health). Mice were recovered on a heat pad, then housed individually for three days following surgery and monitored daily until they recovered to baseline weight.

### Torpor induction

Prior to torpor induction, mice were moved from the home cage to a custom built 32 × 42 × 56 cm cage, divided into four quadrants (16 × 21 × 56 cm), into which each mouse was individually placed. This cage was designed to allow up to 4 animals to be monitored simultaneously using a single thermal imaging camera placed directly above. The Perspex separating each quadrant was clear and had ventilation holes at 2cm from the floor height to allow interaction between mice in neighbouring quadrants, while preventing huddling.

A calorie restriction protocol was used to increase the likelihood of torpor induction (see figure 1) (van der Vinne et al., 2018). Mice were weighed and given a single daily meal placed directly onto the floor of the cage at lights off (ZT0) for five consecutive days. The meal consisted of one pellet (2.2g) of feed (EUROdent Diet 22%, irradiated, 5LF5). This provides 8kcal per day, which is approximately 70% of the estimated unrestricted daily intake for a mouse of this size (Subcommittee on Laboratory Animal Nutrition and Board on Agriculture, 1995).

**Figure 1.**
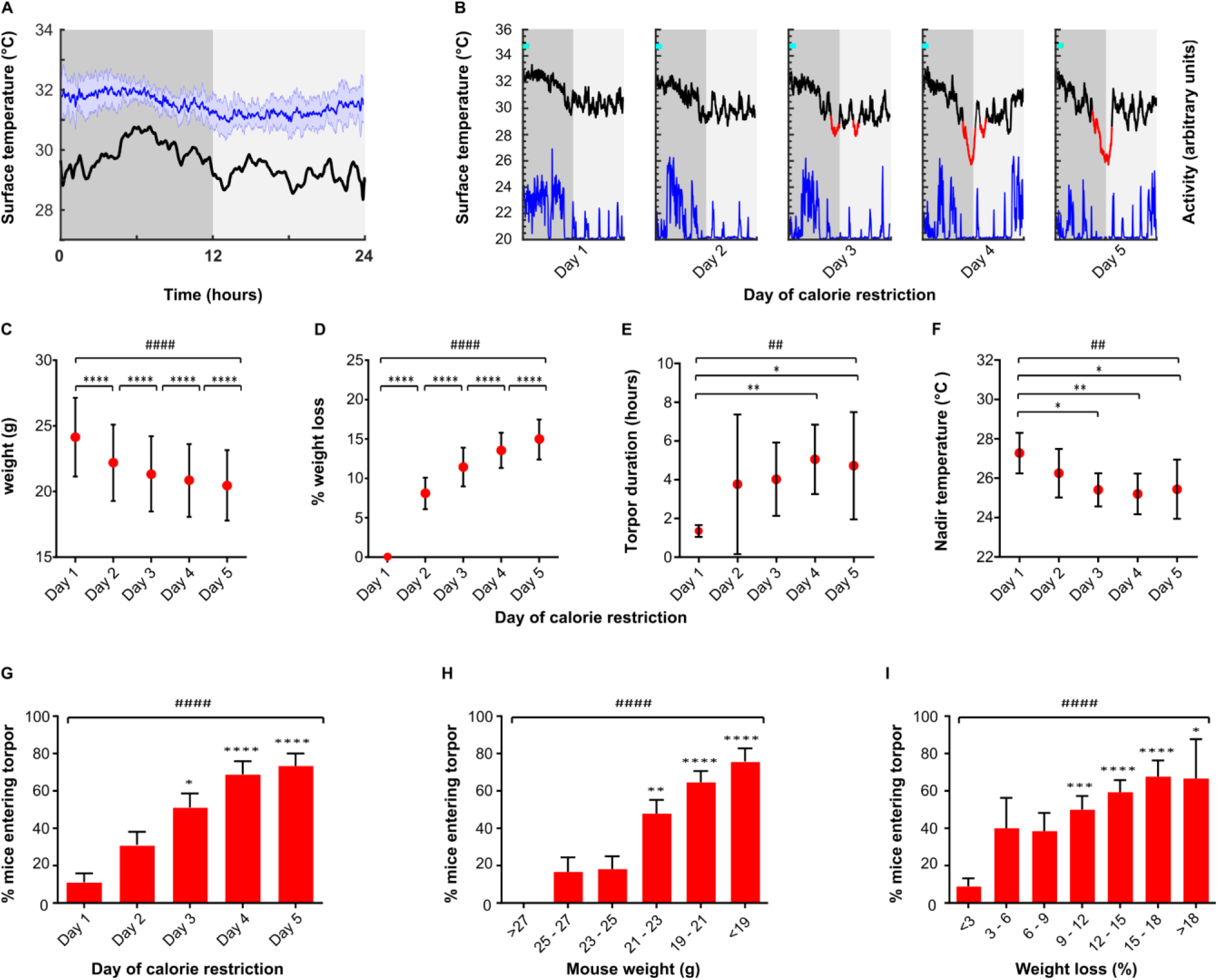
Characteristics of torpor induced by calorie restriction. A, mean mouse surface temperature across the 24-hour day cycle housed at 21°C (n = 7 female mice recorded for three consecutive days, blue line). Shaded area represents standard deviation for that zeitgeber time. Torpor threshold (black line) represents four standard deviations from the mean temperature at that zeitgeber time. B, example profile from single mouse undergoing torpor by 5 days calorie restriction. Mice receive a single meal at lights off (ZT0, cyan marker), providing approximately 70% of the ad-lib daily intake. Surface temperature (black / red line) is measured using infra-red thermography, activity (blue) derived from thermal imaging video. Torpor (red line) defined as a period during which surface temperature remained 4 standard deviations below the mean for that time of day for a period of at least one hour. C & D, Weight loss by day of calorie restriction. E & F, torpor bout duration and nadir surface temperature by day of calorie restriction. G, Proportion of mice entering torpor for each consecutive day of calorie restriction. H, proportion of mice entering torpor by absolute weight. H, proportion of mice entering torpor by % weight loss from first day of calorie restriction. Tests used were Kruskal-Wallis for continuous variables and Friedman test for binary variables. #,##,###,#### = Main effect for day of calorie restriction or weight loss at p <0.05, <0.01, <0.001, or <0.0001, respectively. *,**,***,**** indicate p<0.05, <0.01, <0.001, or <0.0001 respectively for individual comparisons. G-I individual comparisons are relative to mice on day 1. n = 45 trials in 30 female mice, five consecutive days calorie restriction at 21°C ambient temperature with no chemogenetic manipulation.

Mouse surface temperature was recorded using an infra-red thermal imaging camera placed above the cage (Flir C2, www.flir.co.uk). Baseline recordings of mouse surface temperature were taken for a period of three days at an ambient temperature of 21°C. During this time, mice had free access to food and were therefore not expected to enter torpor. These measurements were then used to generate a daily profile mean and a standard deviation of the temperature fluctuations across the diurnal cycle (in one-minute bins).

### TRAP:Ai14 tagging

Female TRAP:Ai14 mice were entered into the calorie restriction protocol and were habituated to daily vehicle injection (chen oil, see below) at ZT7, before receiving 4-OHT on day four, again at ZT7. Approximately half the animals entered torpor within three hours of 4-OHT (50mg/kg), but those that did not were used as controls.

### Chemogenetic targeting of DMH neurons

Adeno-associated Viral vectors were used to deliver either Cre-dependent excitatory (hM3Dq) or inhibitory (hM4Di) DREADD transgenes into the DMH of TRAP2 mice as described above. After at least two weeks recovery, mice were entered into the calorie restriction torpor induction protocol during which they were habituated to daily vehicle injection (chen oil, see below) at ZT7 before receiving 4-OHT on day five, again at ZT7.

Following a further two weeks to allow return to baseline weight and to allow expression of the DREADD protein, mice entered a randomised, crossover design, calorie restriction trial. During this trial, each mouse was randomly assigned to receive either daily CNO (5mg/kg, i.p) or daily saline injections (0.9%, 5ml/kg, i.p) at ZT7, on each of the five days of calorie restriction. The occurrence and depth of torpor was monitored with surface thermography. Following this first arm of the study, and after at least five days with free access to food, the process was repeated with mice that initially received CNO now receiving saline, and vice versa.

To control for a potential influence of CNO on torpor, wild type mice (without a vector injection) underwent the same CNO vs saline crossover trial protocol. To control for an influence of basal DMH activity on torpor, further TRAP2 mice underwent DMH vector injection followed by 4-OHT injection at ZT7 in the home cage with free access to food. These mice were then entered into the calorie restriction protocol with CNO/saline dosing. Torpor duration, and nadir temperature reached were recorded daily for each animal.

Mouse surface temperature was identified by extracting the maximum temperature value from regions of interest within each frame that correspond to each individual mouse’s compartment within the cage. Peak temperature data was extracted from the infra-red video using Flir ResearchIR software version 4.40.9 (www.flir.co.uk), and further filtered and analysed using Matlab 2019a (www.mathworks.com). The data processing stream was as follows: in order to limit noise and movement artefact, data points below 20 or above 40°C were removed; data was then interpolated and resampled at 1 Hz, using the Matlab *interpl* function; finally, a moving average filter function was applied with a 360 data point window, using the Matlab *smooth* function. Mouse activity data was derived from the thermal imaging video, extracted using Ethovision XT software (www.noldus.com).

### Histology: TRAP:Ai14 tagging

Mice were culled by terminal anaesthesia (pentobarbitone (Euthatal) 175mg/kg,) a minimum of four weeks from the time of 4-OHT injection, to allow expression of tdTomato. They were trans-cardially perfused with 10ml heparinised 0.9% NaCl (50 units/millilitre) followed by 20ml of 10% neutral buffered formalin. Brains were removed and stored in fixative solution for 24 hours at 4°C before being transferred to 20% sucrose in 0.1M phosphate buffer (PB), pH 7.4, and again stored at 4°C. Brains were sectioned at 40μm thickness into a 1:3 series on a freezing microtome, transferred to 0.1M PB containing 1:1000 sodium azide. Sections were imaged using a Leica DMI6000 widefield microscope and 0.75 numerical aperture, 20-times magnification objective, excitation filter 546/10nm, dichroic mirror 560nm, emission filter 580/40nm.

Masks for regions of interest were taken from the Mouse Brain Atlas (Franklin and Paxinos, 2007). These masks were then digitally applied to each of the widefield images so that the same size and shape area of interest was used for cell counting across animals. The DMH, PH, and Arc were defined by their atlas boundaries. The preoptic area mask was defined dorsally by the anterior commissure, ventrally by the ventral surface of the brain, and laterally by the lateral extent of the ventrolateral preoptic nucleus. Hence, the preoptic area as defined here included the medial and lateral parts of the medial preoptic nucleus, the ventromedial and ventrolateral preoptic nuclei, the medial and lateral preoptic areas, and parts of the strio-hypothalamic, the septo-hypothalamic, the median preoptic, and the periventricular nuclei (Franklin and Paxinos, 2007). An automated image processing pipeline was employed to count c-fos positive nuclei. This involved background subtraction with a rolling ball radius of 50 pixels. Labelled cells were identified by applying a threshold to the image that identified regions with brightness three standard deviations greater than the mean background. Overlapping regions were separated using the watershed method. Highlighted areas were then filtered by size (50-2000 μm^2^) and counted in Image-J.

### Histology: multiplex RNA in-situ hybridisation

Mice were culled by terminal anaesthesia with intraperitoneal pentobarbitone (175mg/kg, Euthatal). Fresh frozen tissue was prepared, and 15μm coronal sections were cut using a cryostat.

Every third section was taken for standard immunohistochemistry (IHC) to confirm appropriate injection targeting and expression of the DREADD-mCherry fusion protein using a rabbit anti-mCherry primary (Biovision 5993, 1:2000), and donkey anti-rabbit secondaries (Alexafluor594, 1:1000). Sections were imaged using a Zeiss Axioskop II inverted microscope with a CooLED pE-100 excitation system, excitation filter 546/12nm, dichroic mirror 580nm, emission filter 590nm.

For RNAscope (Acdbio, Ca, USA), mounted sections were fixed in fresh 10% neutral buffered formalin solution for one hour at room temperature, then dehydrated using a sequence of increasing concentrations of ethanol (50 – 100%). One slide containing the dorsomedial hypothalamus at bregma −1.94mm was selected from each animal to be processed for multiplex RNAscope for a total of 12 probes (see table 1). Dehydrated sections were air dried for five minutes at room temperature. Slides were then treated with a protease (Protease IV) for thirty minutes at room temperature. The 12 RNA probes were then applied to each slide and hybridised for 2 hours at 40°C in a humidified chamber. After hybridisation, slides were washed in wash buffer twice for 2 minutes at room temperature. Amp 1 was applied to each slide and incubated in a humidified oven at 40°C for thirty minutes, then again washed twice. The process was repeated for two further amplification steps using Amp 2, then 3.

**Table 1.** RNA target probes used across the 12 rounds of ISH. Three RNA targets were imaged in each round, alongside DAPI. The fluorophores were cleaved and then attached to the next round of targets across four rounds. Abbreviations: NPY, neuropeptide Y; TRH, thyrotropin releasing hormone; ChAT, choline acetyl transferase; VGAT, vesicular GABA transporter; VGLUT2, vesicular glutamate transporter 2.

| Fluorophore label | Round 1 | Round 2 | Round 3 | Round 4 |
| --- | --- | --- | --- | --- |
| **AF488** | Leptin Receptor B<br>(cat # 402731) | NPY<br>(cat # 313321) | ChAT<br>(cat # 408731) | mCherry<br>(cat # 431201) |
| **Atto550** | Kappa opioid receptor<br>(cat # 316111) | TRH<br>(cat # 4368110) | Galanin<br>(cat # 400961) | VGLUT2<br>(cat # 319171) |
| **Atto647** | NPY receptor 1<br>(cat # 4270210) | Orexin<br>(cat # 490461) | VGAT<br>(cat # 319191) | NeuN<br>(cat # 313311) |

Next, HiPlex Fluorophore (targeting RNA probes 1-3) was added and incubated in a humidified oven at 40°C for fifteen minutes, then again washed twice for two minutes each time at room temperature. Finally, DAPI was applied to the sections for thirty seconds at room temperature, followed by Prolong Gold antifade mountant (www.thermofisher.com), and a coverslip was applied. Sections were then imaged using a confocal microscope (see below).

After imaging the DAPI, AF488, Atto550, and Atto647 channels, coverslips were removed by soaking in 4X saline sodium citrate (SSC, www.thermofisher.com) for thirty minutes. Cleaving solution (www.acdbio.com) was applied to each section and incubated at room temperature for fifteen minutes to remove the fluorophores from the RNA probe amplifiers, followed by two washes in PBST (PBS with 0.5% Tween). This cleaving process was repeated once.

The entire protocol, from the point of applying the fluorophores to imaging and cleaving, was then repeated three further times, with fluorophores targeting probes 4-6, 7-9, then 10-12 on each successive round). On each occasion DAPI plus three RNA target probes were imaged until all twelve targets had been processed.

Sections for RNAscope were imaged using a Leica SP8 AOBS confocal laser scanning microscope attached to a Leica DMi8 inverted epifluorescence microscope using HyD detectors (‘hybrid’ SMD GaAsP detectors) with 405nm diode and white light lasers. A 40x oil immersion lens with NA of 1.3 (Leica HC PLAPO CS2 lens) was used. Laser settings are shown in Table 1 2.

The images generated from each round were z-projection compressed in Image-J (Schindelin et al., 2012), then proprietary software (www.acdbio.com) was used to align the images from each round for each animal, based on the DAPI staining. Aligned images were then processed in Image-J first applying a 50-pixel radius rolling ball background subtraction. Then, for each fluorescence channel imaged a total of four times for each section (each time labelling a different RNA target probe), a cross session median projection was generated and subtracted from the images generated in each individual round, to subtract any background fluorescence signal.

Each DAPI-stained nucleus was identified and marked with a 12μm diameter area of interest. This created a map of all the nuclei within the imaged section, which could then be projected onto the fluorescence images from each round of RNA target visualisation for that section. DAPI-stained nuclei lining the third ventricle were excluded as these were assumed to represent ependymal cells. Where two DAPI nuclei appeared to overlap, both were excluded on the basis that it would not be possible to distinguish to which nucleus a given RNA probe signal was attached. Nuclei expressing each RNA target were identified manually. From each section, the region immediately lateral to the dorsal part of the third ventricle at bregma −1.94mm was imaged. This included the dorsal and compact parts of the dorsomedial hypothalamus. The area scanned measured 387 × 387 × 10μm (Figure 8).

### Drug preparation

#### Vehicle

The vehicle for 4-hydroxytamoxifen was chen oil, composed of four parts sunflower seed oil and one part castor oil.

#### 4-Hydroxytamoxifen

The z-isomer of 4-hydroxytamoxifen (4-OHT) is the active isomer (www.tocris.com). It was dissolved in chen oil using the following method (Guenthner et al., 2013). First, 4-OHT was dissolved in neat ethanol at 20mg/ml by shaking at 400rpm and 37°C for 30-60 minutes until fully dissolved. Two parts chen oil for every one-part ethanol was then added, and the ethanol was evaporated off using a vacuum centrifuge leaving a final solution of 10mg/ml in chen oil. Drug was prepared on the day of use, and if not used immediately, was kept in solution in the oil by shaking at 400 rpm at 37°C. Once in solution, the drug was protected from light.

#### Clozapine-N-Oxide

CNO was dissolved at 1mg/ml in sterile water at room temperature. Aliquots were stored protected from light for up to one week at room temperature.

#### Experimental design and statistical analyses

During baseline recordings in fed mice (n = 7 female mice recorded for 3 consecutive days), we found that surface temperature frequently dropped below 3 standard deviations from the mean, but infrequently dropped below 4 standard deviations from the mean (see figure 1A and B). Adding a time criterion of 60 minutes duration for the period spent outside of the normal thermoregulatory range optimised the specificity for torpor detection and helped distinguish torpor from sleep: no ad-lib fed mice remained 4 standard deviations below the mean for this length of time, yet visually obvious torpor bouts in calorie-restricted mice were reliably detected. Hence, we defined torpor as any period during which mouse surface temperature remained 4 standard deviations below the mean for that time of day for a period of at least one hour.

Data were analysed using GraphPad Prism (version 6.07). To assay chemogenetic promotion or inhibition of torpor, data from calorie restriction periods in which mice received CNO were compared with data from calorie restriction periods in which the same mice received saline. Data for daily time spent in torpor and nadir temperature were analysed using two-way repeated measures ANOVA with Holm-Sidak’s multiple comparisons test. Analysis of the probability of torpor during calorie restriction used Friedman or Kruskal-Wallis test with Dunn’s multiple comparisons. Comparison of the total number of days with torpor, day of first torpor entry, and weight at first torpor entry when calorie-restricted mice received CNO versus saline were analysed using paired samples *t* test or Wilcoxon matched pairs signed rank test. Normality was assessed by Kolmogorov-Smirnov test. Data is presented as mean ± standard deviation for normally distributed data, or else median [interquartile range]. Where the day of torpor emergence was analysed, if no bout appeared by the end of five days’ calorie restriction, for analyses torpor was assumed to occur on day six.

## Results

### Torpor induction by calorie restriction

Prior to chemogenetic manipulation of torpor-TRAPed neurons, the calorie restriction protocol generated torpor bouts on at least one day in 97% of 45 trials in 30 mice. Mean nadir surface temperature during torpor was 25.3± 1.3°C compared to a mean nadir of 30.0 ± 0.7°C in mice held at the same ambient temperature (21°C) with free access to food (*t*(11) = 9.40, *p* < 0.0001). Entry into torpor occurred in the second half of the lights off period, with the median time of entry into torpor occurring at ZT 9.76 [8.18 – 10.83] hours after lights off.

Torpor duration increased from median 1.27 hours [1.09 – 1.65] on day one to 4.47 hours [1.90 – 6.77] on day five (Kruskal-Wallis test, *H*(5) = 18.05, *p* < 0.01) (see figure 1E). The increase in torpor duration was associated with torpor occurring increasingly early in the day, from a median 12.42 hours [11.21 – 15.13] from lights off on day one to median 9.11 hours [7.87 – 10.77] on day five (Kruskal-Wallis *H*(5) = 20.20, *p* < 0.001). The nadir temperature reached during torpor decreased with increasing days of calorie restriction from 27.3 ± 1.0°C on day one to 25. 4 ± 1.5°C on day five (one-way ANOVA *F*(4,102) = 4.717, *p* <0.01) (see figure 1F). Activity of the mice, derived from the thermal imaging video, reduced to a minimum during torpor, and increased during arousal.

Mice lost weight across the five days of calorie restriction (see figure 1 C & D). Torpor typically emerged on day 3 [2-4], the probability of torpor increased with each day of calorie restriction, with 11.1% of trials resulting in torpor on day 1 (95% CI 3.7 to 24.1%) and 71.1% of trials resulting in torpor on day 5 (95% CI 55.7 to 83.6%) (see figure 1G – I).

### Torpor is associated with increased neural activity in the POA and DMH

To identify neuronal populations that were active during torpor, TRAP:Ai14 mice were calorie-restricted and administered 4-OHT (50mg/kg i.p) on the third day at ZT7 in order to ‘TRAP’ active neurons, with an approximately 50% chance of each mouse entering torpor immediately after 4-OHT injection (n = 5 female mice). We observed widespread recombination resulting in tdTomato expression throughout the hypothalamus, both in mice that entered torpor (Torp+) following 4-OHT, and those that did not (Torp-, see figure 2).

**Figure 2.**
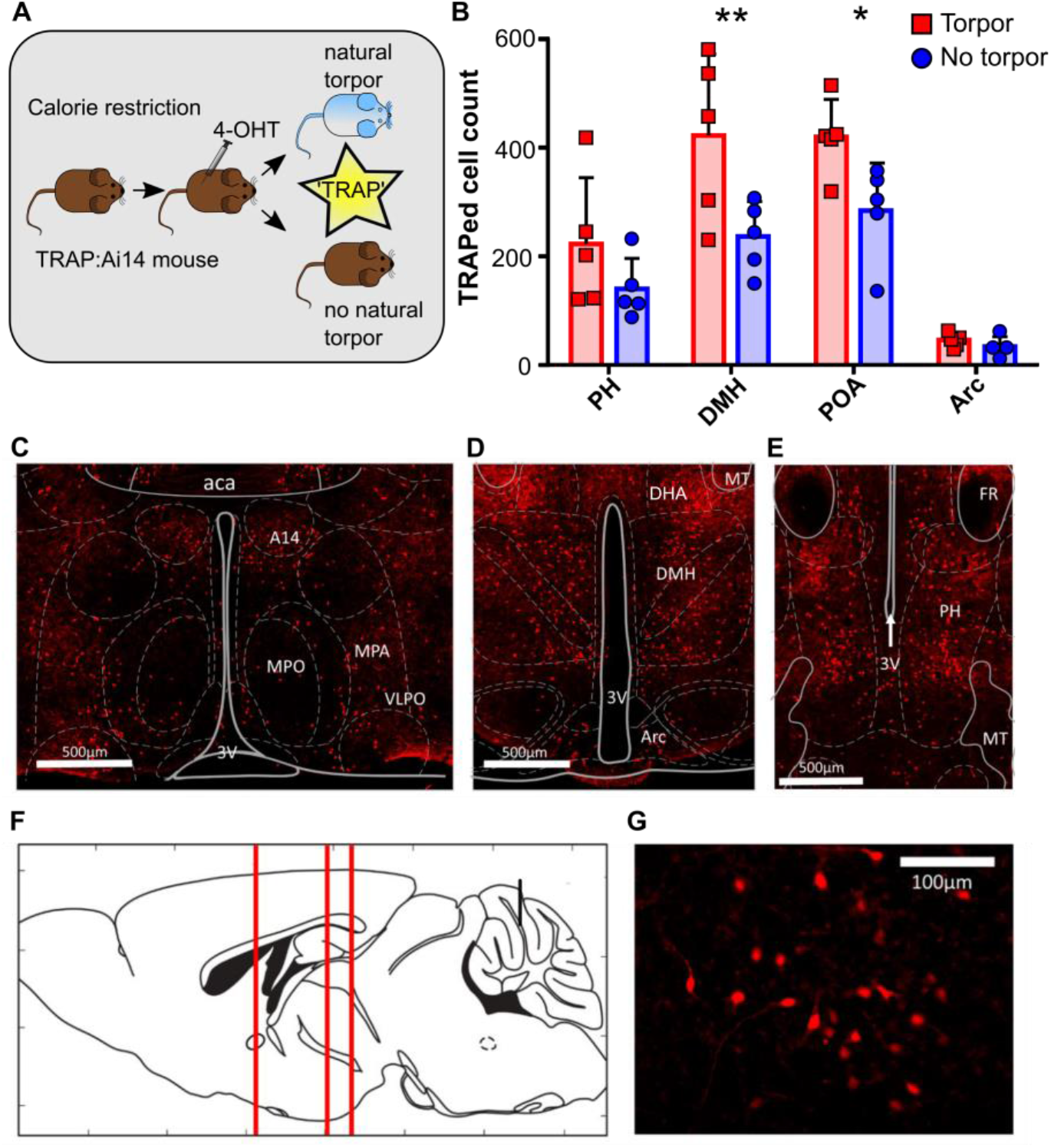
Torpor is associated with increased activity in dorsomedial and preoptic hypothalamus (n = 5 female TRAP:Ai14 mice). A, torpor TRAP protocol for TRAP2:Ai14 mice. Calorie restricted TRAP:Ai14 mice receive 4-hydroxytamoxifen (4-OHT) at ZT7, those that entered torpor following the injection are compared with those that did not. B, Counts of td-Tomato-positive cells by region of interest in calorie-restricted mice that entered torpor following 4-OHT (red) or did not enter torpor following 4-OHT (blue). Torpor was associated with higher numbers of ‘TRAPed’ cells in the DMH and POA compared to animals that did not enter torpor. RM ANOVA comparing TRAPed cell count in mice that entered torpor with those that did not for each of the four regions of interest. Main effect for torpor vs no torpor, p < 0.05, main effect for region of interest, p < 0.0001, significant torpor vs no torpor by region of interest interaction, *p* < 0.05. Holm-Sidak’s multiple comparisons test,** indicates *p*<0.01, * indicates *p*<0.05. Data shown are mean and standard deviation. C, coronal section showing POA neurons; D, DMH and Arc neurons; E, PH neurons. F, sagittal schematic showing corresponding anterior-posterior location of coronal sections C, D, and E; G, high magnification image showing DMH torpor-active neurons expressing td-Tomato. Abbreviations: aca, anterior commissure (anterior part); A14, A14 dopamine cells; Arc, arcuate nucleus; DHA, dorsal hypothalamic area; DMH, dorsomedial hypothalamus; MPO, medial preoptic nucleus; MPA, medial preoptic area; PH, posterior hypothalamic nucleus; POA, preoptic area; VLPO, ventrolateral preoptic nucleus; 3V, third ventricle. n = 5 female mice per group.

Four hypothalamic areas were selected *a-priori* for cell counting: POA, DMH, posterior hypothalamus (PH), and arcuate nucleus (arc). Entry into torpor was associated with increased tdTomato labelled (‘TRAPed’) cells in the DMH and POA as compared to calorie-restricted mice that did not enter torpor (DMH Torp+ versus Torp-mean difference 185.6 TRAPed neurons, confidence interval (CI) 76 – 295, POA mean difference 135 TRAPed neurons, CI 26 – 245, Holm-Sidak multiple comparisons test *p* < 0.01 and *p* < 0.05, respectively). In contrast, the arcuate and posterior hypothalamus did not show significant differences between Torp+ and Torp-mice (see figure 2B). These differences in activity in the POA and DMH were not due to greater weight loss in the mice that entered torpor, indicating that the increased activity related to the occurrence of torpor *per se* rather than a response to the degree of calorie deficit (mean weight loss Torp+ 2.6 ± 0.7g, versus 2.4 ± 0.3g Torp-, *t*(8) = 0.66, *p* = 0.46; percentage weight loss Torp+11.6 ±2.3% versus 11.0 ± 1.2% Torp-, *t*(8) = 0.57, *p* = 0.58).

### Chemoactivation of torpor-TRAPed DMH neurons

To test whether neurons within the DMH play a causal role in torpor, we targeted expression of the excitatory DREADD hM3Dq to neurons in the DMH that were active during torpor. Nine TRAP2 mice had injection of the Cre-dependent excitatory DREADD AAV (AAV2-hSyn-DIO-hM3Dq-mCherry) into the dorsomedial hypothalamus, followed by five days of calorie restriction with 4-OHT injection on day five (see figure 3A). Of these mice, seven entered torpor following 4-OHT administration and were included in subsequent experiments (Torp+ hM3Dq).

**Figure 3.**
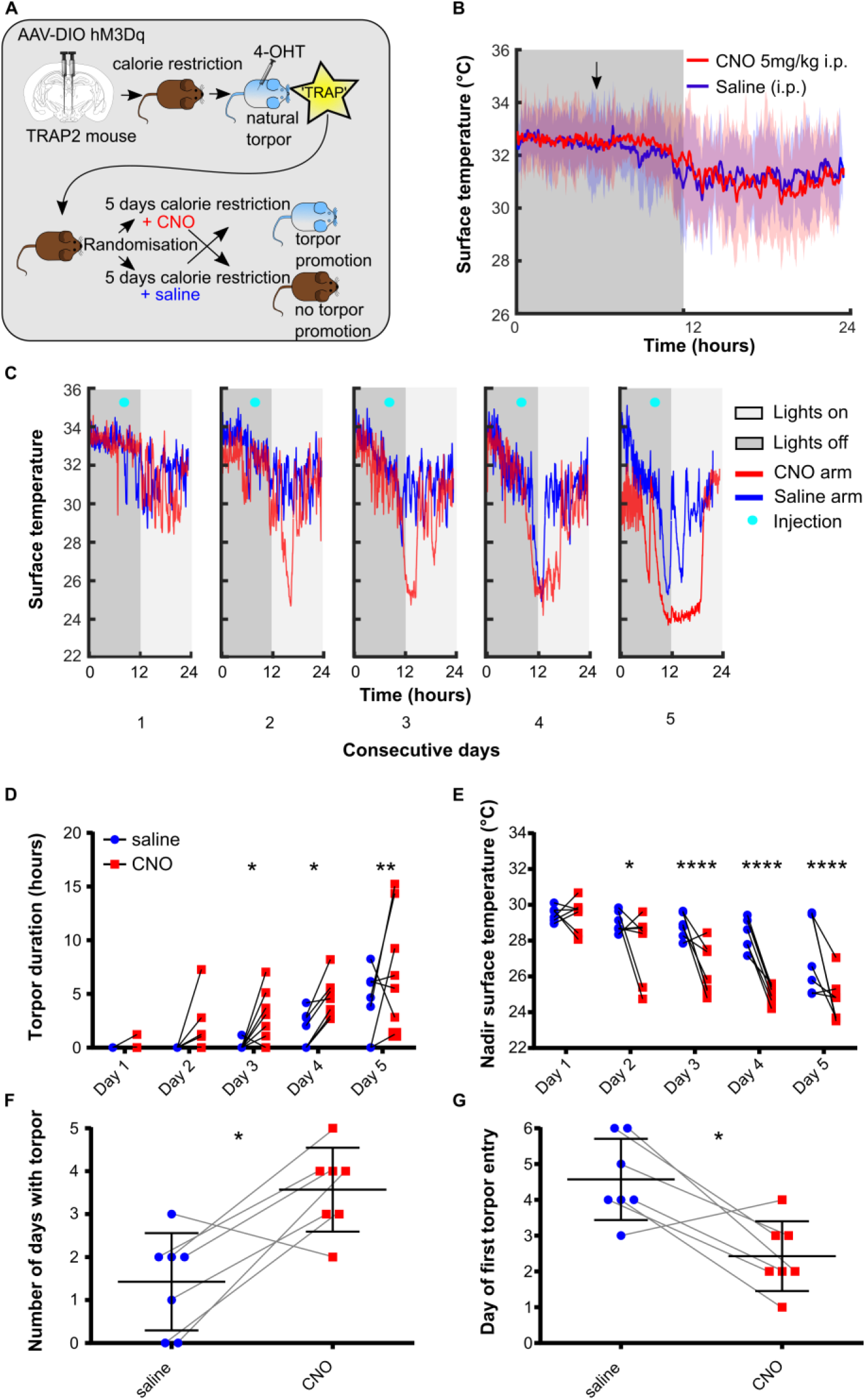
Chemoactivating DMH torpor-TRAPed neurons promotes, prolongs, and deepens torpor bouts in calorie restricted mice (n= 7 female Torp+ hM3Dq mice). A, DMH injection and torpor-TRAP protocol. B, CNO does not trigger torpor in DMH torpor-TRAPed mice that are not calorie restricted (shaded area represents 95% CI). C, example plot showing surface temperature in calorie restricted DMH torpor-TRAPed mice receiving CNO (red) or saline (blue) at 7 hours after lights off (cyan marker). D, CNO given to calorie restricted DMH torpor-TRAPed mice increases the total time spent in torpor and decreases the nadir surface temperature reached (D & E, respectively. 2-way repeated measures ANOVA, significant main effect for CNO vs saline trials, *p* < 0.05 for both torpor duration and nadir temperature. Holm-Sidak’s multiple comparisons test *,**,***,***, indicate significant difference between CNO and saline on individual days at *p* < 0.05, 0.01, 0.001, 0.0001, respectively). CNO increased the total number of days in which torpor occurred and resulted in torpor occurring after fewer days of calorie restriction (F & G, respectively. * Indicates Wilcoxon matched-pairs signed rank test or paired *t*-test, *p* < 0.05, comparing number of torpor bouts and day of first torpor bout when mice received CNO versus saline).

**Figure 4.**
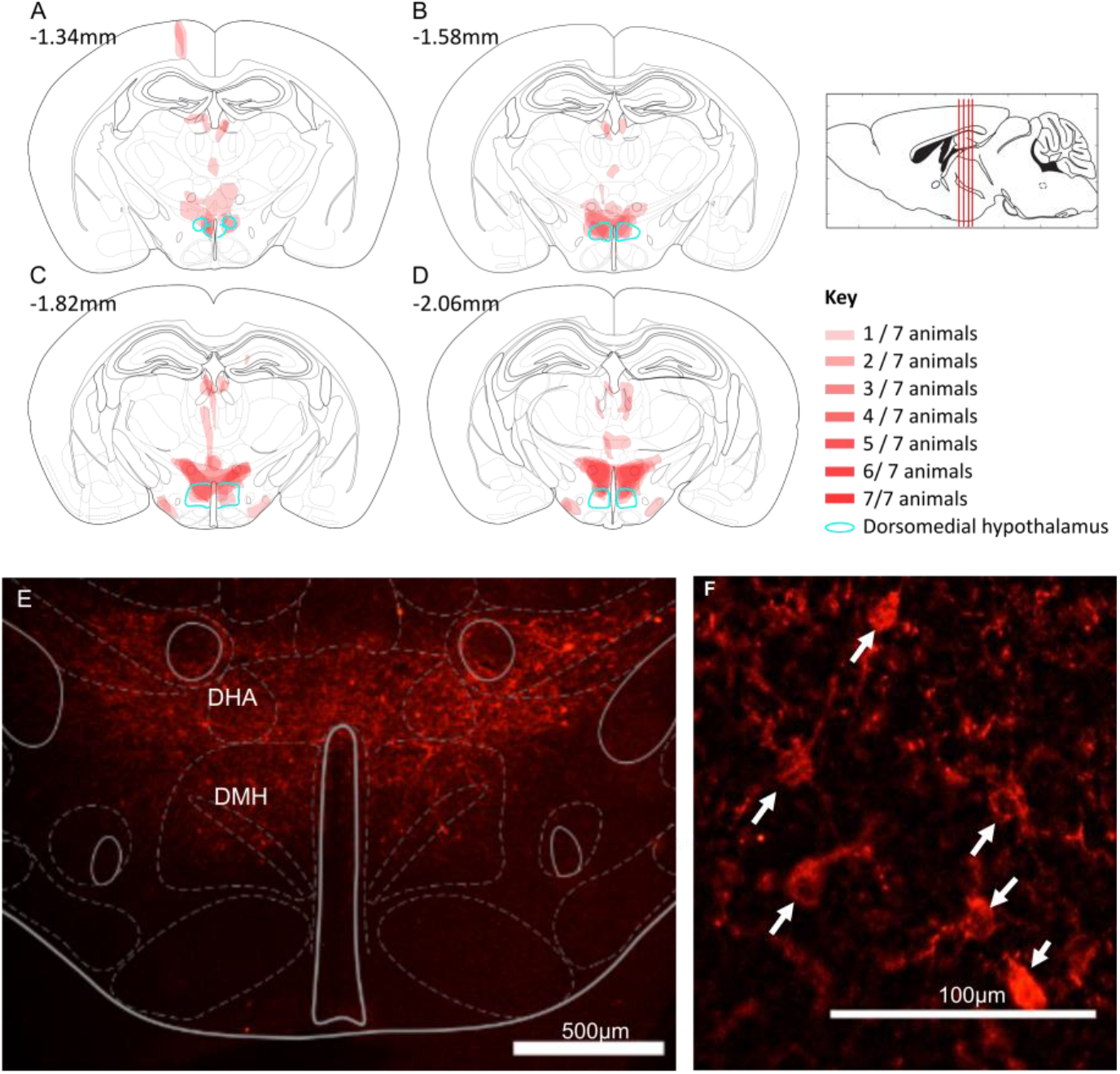
Torpor-TRAPed DMH neurons (n = 7 female Trap+ hM3Dq mice). A-D, Mapped extent of mCherry-labelled cell bodies, indicating TRAPed cells expressing hM3DGq. TRAPed cells were observed in the dorsomedial hypothalamus (DMH, marked in green) and dorsal hypothalamic area (DHA) of all mice. Injection tracts visible in A at bregma −1.34mm, resulting in TRAPed cells in the cortex. Two mice also showed mCherry labelled cell bodies in the medial tuberal nucleus (visible in C & D). E & F, example mCherry labelling (red) indicating DREADD expression in TRAPed cells in an hM3Dq-DMH-TRAP mouse. Expression in the dorsomedial hypothalamus (DMH) and dorsal hypothalamic area (DHA). Immunohistochemistry performed labelling the mCherry component of the hM3Dq-mCherry fusion protein. F, Arrows indicate mCherry labelled neuron cell bodies. 15μm coronal sections.

Chemoactivation of DMH torpor-TRAPed neurons by intraperitoneal injection of CNO to fed mice did not trigger torpor, nor did it alter surface temperature (see figure 3B). However, CNO delivered each day at ZT7, to calorie restricted mice, increased the probability of the mice entering torpor over five days (see figure 3C-G). Torpor emerged after fewer days of calorie restriction when mice received CNO compared to when the same mice were calorie restricted and received saline (first torpor bout appeared on day 2.4±1.0 (CNO) versus 4.6±1.1 (saline), paired *t*(6) = 3.60, *p* < 0.05). There were more torpor bouts per mouse on the CNO arms compared to the saline arms (3.6±1.0 (CNO) versus 1.4±1.1 (saline) bouts, paired t(6) = 3.60, *p* < 0.05). Likewise, when mice received CNO during the five days of calorie restriction, torpor bouts were longer than in the arms when they received saline (2-way repeated measures ANOVA main effect for CNO versus saline, *F*(1,6) = 8.14, *p* < 0.05). Similarly, during calorie restriction trials in which mice were given CNO, the daily nadir temperature reached was lower than when they were given saline (2-way repeated measures ANOVA main effect for CNO versus saline, *F*(1,6) = 12.2, *p* < 0.05).

The weights of mice on entry into the CNO vs saline arm of the trial were not different (24.7 ±2.7g (CNO) versus 24.7 ±2.7g (saline), paired *t*(6) = 0.049, *p* = 0.96). Hence, the increased propensity to enter torpor observed with chemoactivation of hM3Dq-DMH-torpor-TRAP mice was not due to systematic differences in their weights.

### Chemoinhibition of torpor-TRAPed DMH neurons

Twelve TRAP2 mice underwent injection of the Cre-dependent inhibitory DREADD (pAAV2-hSyn-DIO-hM4Di-mCherry) into the dorsomedial hypothalamus followed by five days of calorie restriction with 4-OHT injection on day five. Of these mice, six entered torpor following 4-OHT administration and were included (Torp+ hM4Di). These were evaluated for inhibition of torpor in response to CNO compared to saline.

Chemoinhibition of torpor-TRAPed DMH neurons in calorie restricted mice did not reduce the total number of days in which torpor occurred (0.8±1.2 (CNO) versus 1.7±1.2 bouts (saline), paired *t*(6) = 1.05, *p* = 0.34). Nor did it delay the onset of torpor across the five days of calorie restriction (torpor appeared on day 5.2±1.2 (CNO) versus day 4.3 ±1.2 days (saline), paired *t*(5) = 1.05, *p* =0.34) (see figure 5). There were no differences between calorie restriction trials in which mice received CNO compared to when they received saline, in terms of time spent in torpor or nadir surface temperature reached (2-way repeated measures ANOVA main effect for CNO versus saline, *F*(1,5) = 1.37, *p* = 0.29 and *F*(1,5) = 1.06, *p* = 0.35, respectively). Mouse weight on entry into the CNO arms was not different to that in the saline arms of the trial (25.5 ±1.9g (CNO) versus 25.2 ±2.1g (saline), paired *t*(5) = 0.73, *p* = 0.5). Hence, chemoinhibition of Torp+ hM4Di mice did not alter the expression of torpor during five days of calorie restriction.

**Figure 5.**
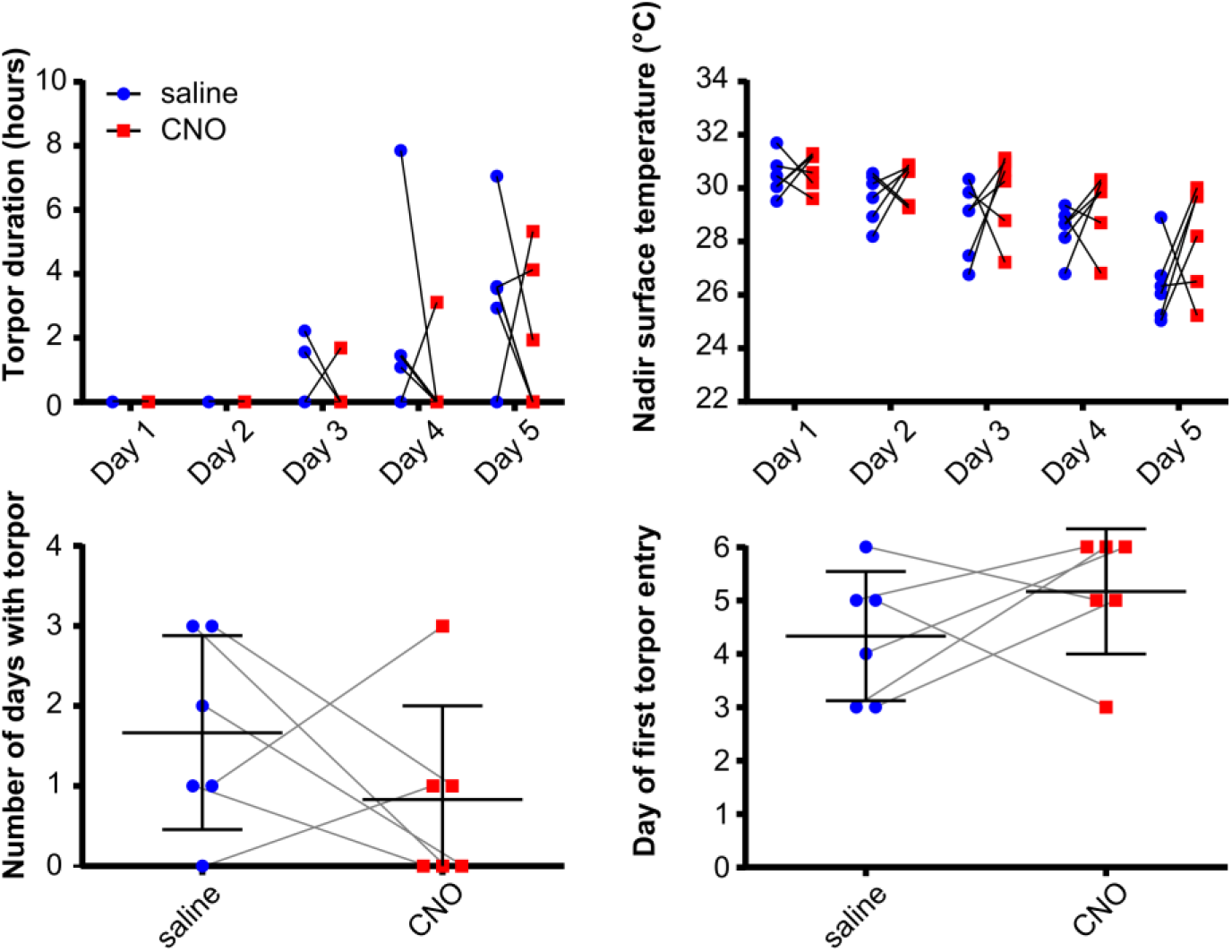
Chemoinhibition of DMH torpor-TRAPed DMH neurons does not affect torpor in calorie restricted mice (n = 6 female Trap+ hM4Di mice). CNO did not affect the duration of torpor bouts, or nadir surface temperature reached (2-way repeated measures ANOVA, no main effect for CNO vs saline across all days, with Fisher’s least significant difference test for CNO versus saline at each day of calorie restriction for each measure, *p* >0.05 throughout). CNO did not affect the total number of days in which torpor occurred, nor the first day on which torpor occurred during five days of calorie restriction (paired *t*-test, *p* > 0.05). n = 6 female mice.

### Controls

Five mice underwent hM3Dq vector injection into the DMH followed by 4-OHT at ZT7 with free access to food in the homecage (Homecage hM3Dq). They were then entered into the randomised, crossover design, calorie restriction trial receiving CNO then saline or vice versa. Again, chemoactivation of DMH neurons TRAPed in the fed homecage condition had no effect on torpor, in terms of the day of first torpor bout appearance (4 [1.5-5.5] CNO, versus 3 [2-3], Wilcoxon matched-pairs signed rank test, W = 6.0, *p* = 0.5, saline), total number of torpor bouts across five days calorie restriction (2.8±1.8 versus 3.2 ±0.84, paired *t*(4) = 0.43, *p* = 0.69), daily nadir surface temperature (2-way repeated measures ANOVA main effect for CNO versus saline, *F* (1, 4) = 0.20, *p* = 0.68), or daily torpor bout duration (2-way repeated measures ANOVA main effect for CNO versus saline, *F* (1, 4) = 0.06, *p* = 0.83) (figure 6).

**Figure 6.**
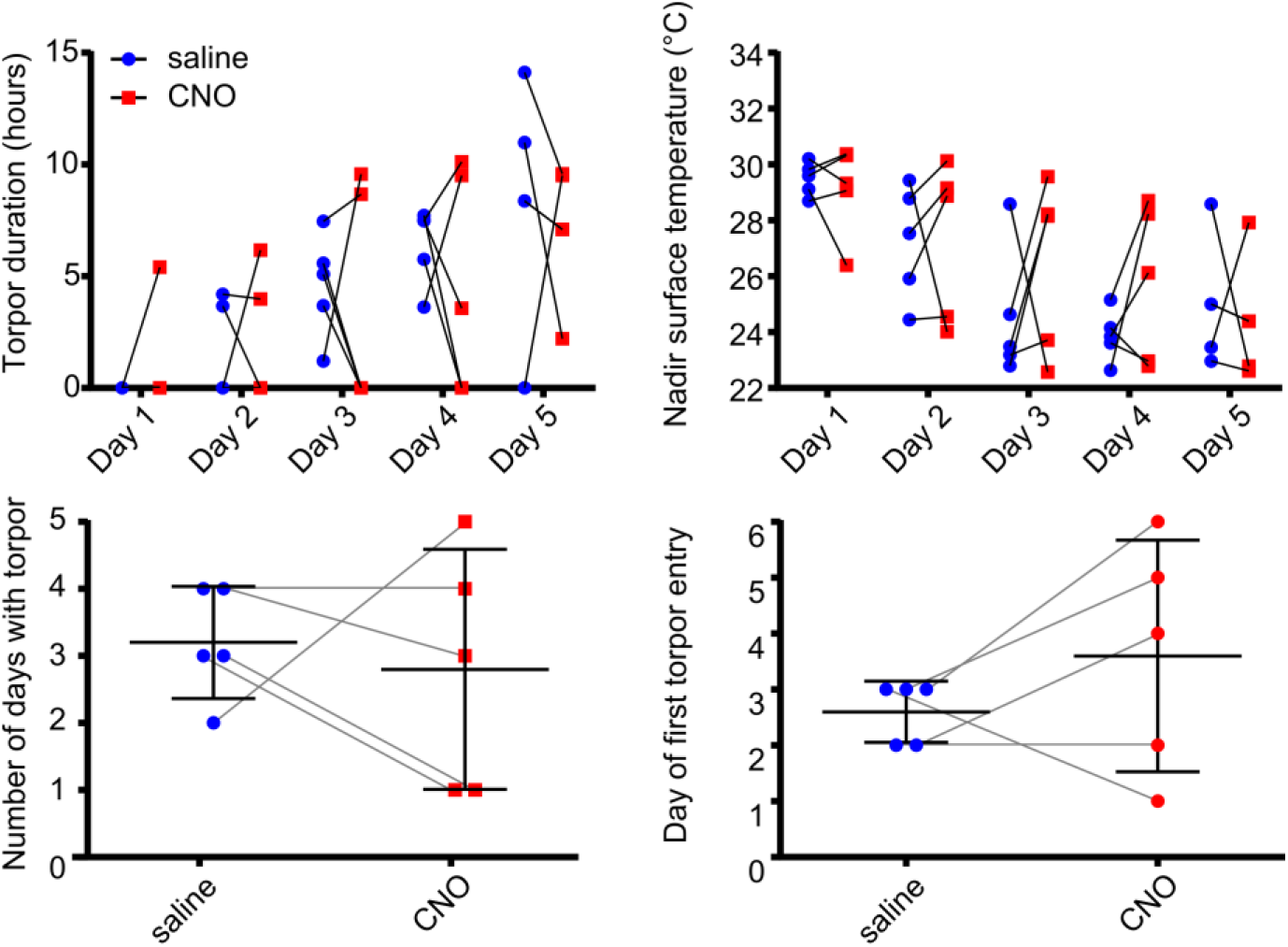
CNO administration does not affect torpor in DMH homecage-TRAPed mice. CNO did not affect the duration of torpor bouts, or the nadir surface temperature reached (2-way repeated measures ANOVA, no main effect for CNO vs saline across all days, with Holm-Sidak’s multiple comparisons for CNO versus saline at each day of calorie restriction for each measure, *p* >0.05 throughout). CNO did not affect the total number of days in which torpor occurred, nor the first day on which torpor occurred during five days of calorie restriction (paired *t*-test, *p* > 0.05).

CNO injection had no effect on torpor in wild type calorie restricted control mice (n=4), in terms of the day of first torpor bout appearance (3 [0.5-4.75] versus 2 [0-4], Wilcoxon matched-pairs signed rank test, W = −3.0, *p* = 0.75), total number of torpor days in which torpor occurred across the five days calorie restriction (3 [1.25-5.5] versus 4 [2-6], Wilcoxon matched-pairs signed rank test, W = 3.0, *p* = 0.75), daily nadir surface temperature (2-way repeated measures ANOVA, *F* (1, 3) = 0.04, *p* = 0.85), or daily torpor bout length (2-way repeated measures ANOVA, *F* (1, 3) = 0.10, *p* = 0.78) (figure 7).

**Figure 7.**
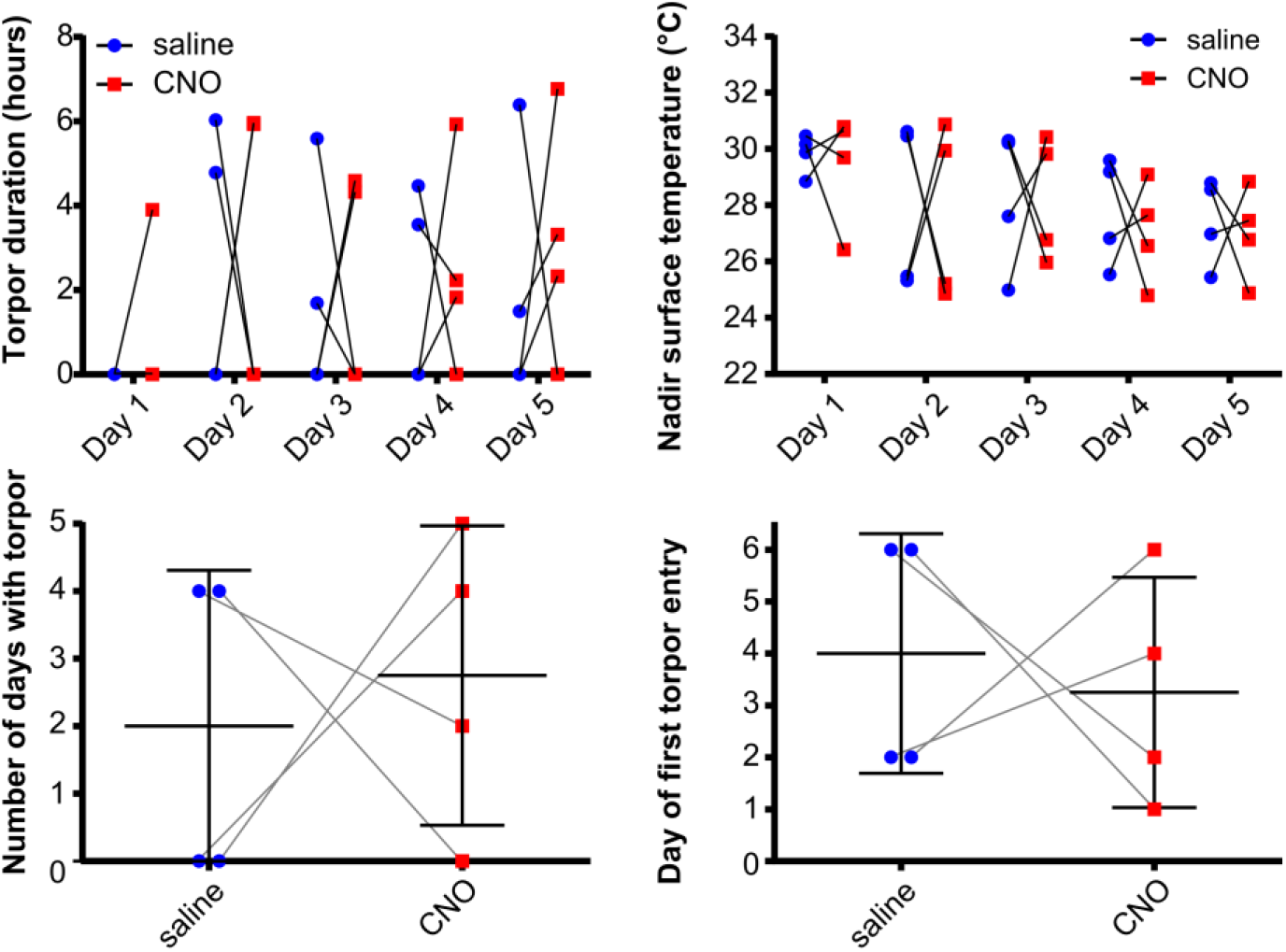
CNO administration does not affect torpor in wild type mice (n = 4 female mice). CNO did not affect the duration of torpor bouts, or the nadir surface temperature reached (2-way repeated measures ANOVA, no main effect for CNO vs saline across all days, with Holm-Sidal’s multiple comparisons test for CNO versus saline at each day of calorie restriction for each measure, *p* >0.05 throughout). CNO did not affect the total number of days in which torpor occurred, nor the first day on which torpor occurred during five days of calorie restriction (paired *t*-test, *p* > 0.05).

### Phenotyping DMH torpor-TRAPed neurons

Sections from Torp+ hM3Dq mice were selected for RNAscope analysis (n = 5 sections from 5 female mice, Figure 8). One target probe, NeuN, did not produce a meaningful pattern of labelling. Because of the failure of NeuN, we were unable to exclude non-neuronal cells from the analysis.

**Figure 8.**
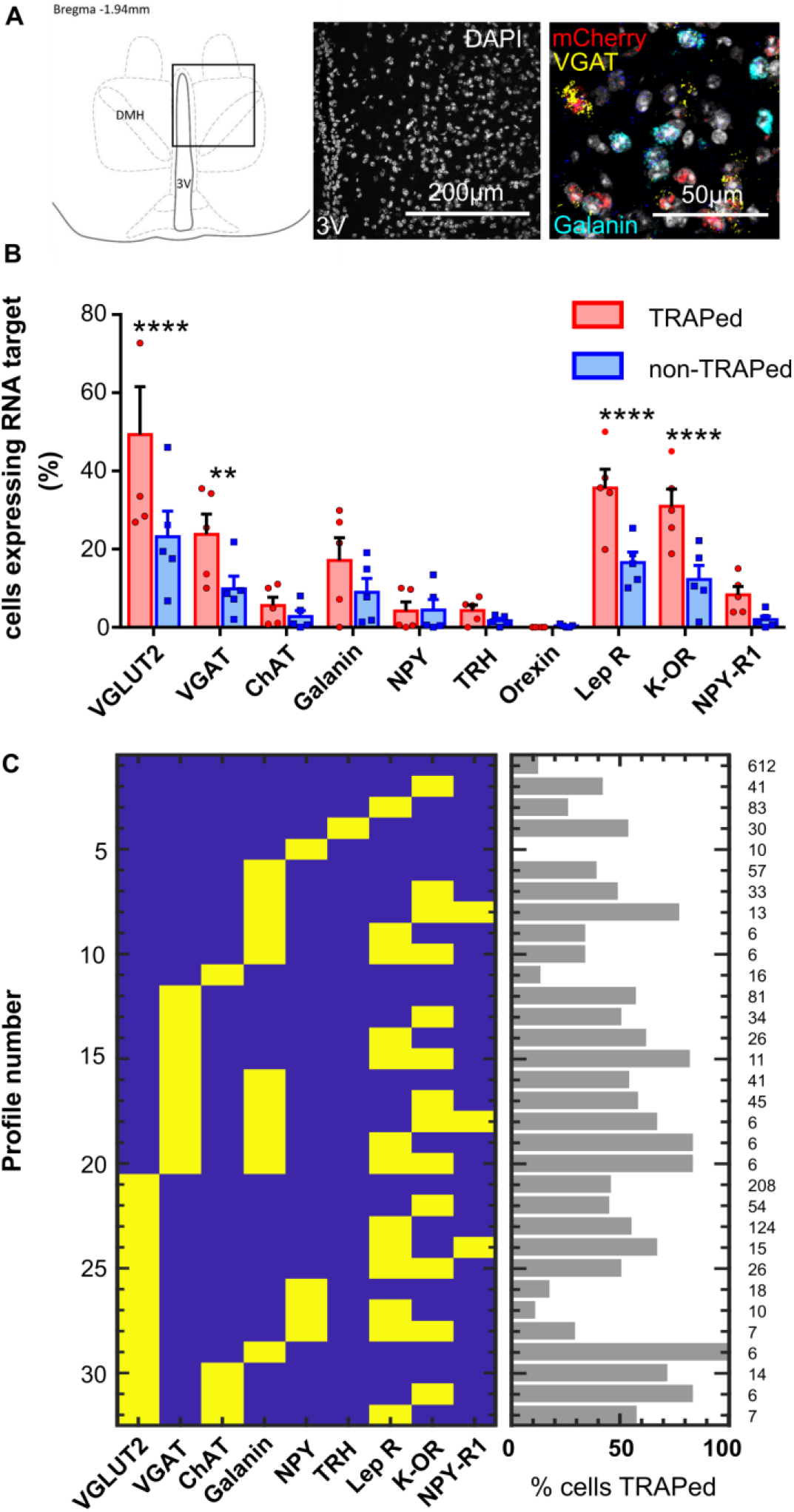
multiplex RNA in-situ hybridisation (ISH, RNAscope) data (n = 5 Torp+ hM3Dq female mice). A, representative section showing DMH region analysed with RNA in-situ hybridisation. B, Proportion of TRAPed versus non-TRAPed cells that expressed each RNA target (2-way ANOVA with Holm-Sidak’s multiple comparisons tests assessing difference in expression of the RNA target in TRAPed compared to non-TRAPed cells ** indicates *p* <0. 01, **** indicates *p* 0.001). C, patterns of RNA target expression observed in the DMH (excluding patterns that occurred in fewer than 5 cells).

A total of 1771 DAPI-stained nuclei were identified across five sections from five animals (345 ±65 cells per animal). Four RNA targets were significantly more likely to be expressed in TRAPed cells compared to non-TRAPed cells: VGLUT2, VGAT, leptin receptor B, and κ-opioid receptor (2-way ANOVA with Holm-Sidak’s multiple comparisons tests, *p* <0.001, *p* < 0.01, *p* 0.001, and *p* < 0.001, respectively, see figure 8). Amongst cells expressing these RNA targets, we found that VGLUT2 and leptin receptor were commonly co-expressed (38% of VGLUT2 cells also expressed leptin receptor compared to 21% that also expressed κ-Opioid R), while VGAT and κ-opioid receptor were commonly co-expressed (38% of VGAT expressing cells also expressed κ-Opioid R, whereas 23% of VGAT cells expressed leptin receptor).

## Discussion

The data presented here support the hypothesis that both the DMH and the POA contain neurons that are active in the period leading up to a torpor bout. We show here that chemoactivation of DMH torpor-TRAPed neurons increases the probability, as well as the depth and the duration of torpor in calorie restricted mice.

The POA has been implicated in the induction of torpor, or torpor-like states in mice (Hrvatin et al., 2020; Takahashi et al., 2020; Zhang et al., 2020). Previous studies have also identified that the DMH contains neurons that express c-fos around the time of torpor (Hitrec et al., 2019; Hrvatin et al., 2020), and several studies have demonstrated that projections from the POA to the DMH can decrease body temperature (Song et al., 2016; Takahashi et al., 2020; Zhao et al., 2017). However, a specific role for the DMH in the control of torpor, rather than a homeostatic thermoregulatory, or counter-regulatory role has not been established. Here, by chemogenetically reactivating specifically those DMH neurons that expressed c-fos around the time of torpor, we have shown that they do indeed contribute to the generation of torpor.

Chemoactivation of DMH torpor-TRAPed neurons did not alter body temperature after single doses of CNO delivered to mice with free access to food. This suggests that the TRAPed neurons play a specific role in promoting torpor under calorie restricted conditions, but that they are not sufficient to trigger it in the fed state. They likely form part of a chorus of signals that indicate negative energy balance and the need to engage torpor. If, on the other hand, the TRAPed neurons were simply part of a thermoregulatory circuit that inhibits thermogenesis under ‘normal’ homeostatic conditions, then one might expect chemoactivation to produce a physiological response that is independent of calorie restriction.

Whether there is a single group of neurons capable of inducing torpor, i.e. a torpor ‘master switch’ remains unknown, although emerging evidence places the POA as a potential candidate (Hrvatin et al., 2020; Takahashi et al., 2020; Zhang et al., 2020). The observation of torpor-promoting effects following chemoactivation of DMH neurons supports the hypothesis that a network of regions contributes to torpor induction. The DMH would then be considered one part of this network. If neurons in the POA are sufficient to trigger torpor, then the DMH might form part of the afferent signal to the POA, or it might modulate the descending efferent signals from the POA.

Another question is whether torpor is triggered, maintained, and terminated by the activity of a single population of neurons, or whether different populations are each responsible for timing the different phases. The observation that chemoactivating DMH torpor-TRAPed neurons increases bout duration as well as increasing the likelihood of torpor occurring hints that torpor may be induced and maintained by the same population of neurons. If, on the other hand, the effect of chemoactivation of DMH torpor-TRAPed neurons was to increase the probability but not the duration of torpor (or vice versa), then this would support the hypothesis that these phases of torpor are governed by distinct neuronal populations.

The finding that activation of neurons with in the DMH contributes to torpor induction - a physiological response characterised by suppression of thermogenesis - is particularly interesting. Within the framework of our current understanding of hypothalamic thermoregulatory circuits, the DMH generally emerges as a driver rather than inhibitor of thermogenesis, at least in rodents (Morrison and Nakamura, 2019; Rezai-Zadeh et al., 2014; Zhao et al., 2017), although cholinergic neurons in the DMH may suppress brown adipose tissue thermogenesis (Jeong et al., 2015). In contrast, the DMH neurons TRAPed during torpor in this study appear to contribute to the suppression of thermogenesis associated with torpor, supporting a more complex and bi-directional role for the DMH in temperature regulation (Saper and Machado, 2020).

We used RNAscope to explore the phenotype of torpor-TRAPed cells within the DMH. We appear to have predominantly TRAPed two populations of neurons: one expressing VGLUT2 and Leptin Receptor B, and the other expressing VGAT and κ-Opioid R. The VGAT and κ-opioid receptor expressing cells might represent a population of neurons that project to the raphe pallidus to inhibit thermogenesis during torpor (Hitrec et al., 2019). Indeed, central blockade of κ-opioid receptors blocks the hypothermic response to calorie restriction in male mice (Cintron-Colon et al., 2019). The VGLUT2 and leptin receptor expressing neurons are also interesting. A similar population have been described previously, as having a role in thermogenesis and maintenance of stable body weight (Zhang et al., 2011; Rezai-Zadeh et al., 2014). If the glutamatergic/leptin receptor-expressing population we have TRAPed is the same as those previously identified, then these neurons might have a role in thermogenesis during the emergence from torpor. Alternatively, they might represent a novel population of leptin-sensitive neurons with a role in torpor promotion.

Finally, we note that an population of neurons that express genes not targeted in our selection of RNA probes might have been critical to the promotion of torpor presented here. For example, we did not include other VGLUT subtypes, and we found 7% of cells that expressed mCherry expressed no other RNA target. These might be important contributors to the promotion of torpor observed in our experiments. The failure of the RNA NeuN probe in our hands limits interpretation of the cell phenotyping data because non-neuronal cells could not be excluded from the analysis and so these findings should be regarded as preliminary and hypothesis generating for future studies.

### Limitations

We opted to deliver 4-OHT in anticipation of torpor, in an effort to reduce the likelihood of arousing the mice and TRAPing arousal circuits. It is possible that this approach resulted in TRAPing neurons within the DMH that are not involved in torpor induction, and/or missed populations whose activity increased nearer to the point of torpor induction. Hence, it remains possible that delivery of 4-OHT during torpor might TRAP a population of DMH neurons capable of inducing torpor in the absence of calorie restriction.

We did not find that chemoinhibition of DMH torpor-TRAPed neurons prevented torpor induction in calorie restricted mice. It is worth noting that incomplete suppression of neuronal firing with the inhibitory DREADD, hM4Di, has been reported (Smith et al., 2016). Therefore, this result could reflect failure to inhibit these DMH neurons sufficiently to prevent their contribution to torpor. Alternatively, it might be necessary to silence a wider population of neurons, including potentially those in other brain regions, to prevent torpor.

Finally, we used female mice in our experiments, because in our experience they have a greater tendency to enter torpor. Whether our results are generalisable to male mice, and whether the stage of the ovarian cycle interacts with the expression of torpor is currently unknown. Recent data suggests neurons in the POA that express oestrogen receptors may have a role in torpor expression, although the effect on torpor of modulating the activity of these oestrogen receptors has not been established (Zhang et al., 2021, 2020). We do not believe that the delivery of 4-OHT during the TRAPing process influenced the results observed here (and our findings are in line with those of Hrvatin et al who also used 4-OHT (Hrvatin et al., 2020)). 4-OHT is generally considered an oestrogen receptor antagonist (Shiau et al., 1998), and we observed no effects on torpor propensity in control mice that received 4-OHT under fed homecage conditions.

### Conclusions

Our data indicate that neurons within the DMH promote torpor entry and increase the duration and depth of torpor bouts in calorie restricted mice. Chemoactivating these neurons in fed mice neither triggered torpor nor altered thermoregulation. Hence, we hypothesise that the DMH plays a modulatory role capable of promoting torpor. Important future work should clarify the phenotype of these neurons and investigate their interaction with other areas believed to be involved in torpor, such as the POA.

## Acknowledgements

This research was funded in whole, or in part, by the Wellcome Trust 211029/Z/18/Z. For the purpose of open access, the author has applied a CC BY public copyright licence to any Author Accepted Manuscript version arising from this submission. MA was funded by a GW4 CAT Wellcome clinical research training fellowship; TH was funded by the Medical Research Council, UK (MR/P025749/1); MC was funded by the European Space Agency (Research agreement collaboration 4000123556). The authors gratefully acknowledge the Wolfson Bioimaging Facility for their support and assistance in this work.

